# Can we tell where our bodies end and the world begins? Evidence for precise three-dimensional internal body models

**DOI:** 10.1101/2025.03.28.645649

**Authors:** Celia R Blaise, Holly C Clark, Hannes P Saal

**Affiliations:** School of Psychology, University of Sheffield, Sheffield, UK; Insigneo Institute for in silico Medicine, University of Sheffield, Sheffield, UK

**Keywords:** body representation, psychophysics, spatial cognition

## Abstract

Distinguishing our body from the external world is crucial for our sense of self, environmental interaction, and sensory processing. Yet the accuracy with which we perceive this boundary remains underexplored. Here, we developed a novel psychophysical protocol to directly assess how accurately individuals perceive their body boundaries. Participants were asked to determine whether the midpoint between two tactile stimuli applied on the skin was inside or outside their perceived body boundary. 3D scans of the tested regions were used to determine objective anatomical boundaries, allowing psychometric functions to be fitted. Results revealed remarkable overall accuracy, on average within millimeters, in localizing body boundaries across multiple body regions. However, accuracy was not uniform: while palm boundaries were localized with near-perfect precision, stimuli straddling the wrist boundaries were frequently misjudged as extending beyond their true anatomical limit, revealing a systematic perceptual bias. Although perceptual judgments adapted to changes in posture, accuracy declined when the detailed local 3D structure was disregarded, indicating that proprioceptive cues are combined with detailed local body models. Finally, participants whose anatomy deviated from the average tended to align their responses with a typical body model rather than their unique physiology, suggesting that top-down processes influence boundary judgments. Together, our findings suggest that body boundary representation combines detailed three-dimensional body models with proprioceptive feedback into an integrated perceptual model of the anatomical body.

## Introduction

Our sense of where our body ends and the external world begins can fluctuate dramatically. At times, this boundary can feel sharp and distinct, such as when plunging into a cold swimming pool. In other moments, it appears fuzzy or indistinct: reclining on a comfortable couch can blur the sense of where one’s body ends and the couch begins (p. 97 in Ihde, 1973; Ratcliffe, 2013). Importantly, our perception of this boundary does not necessarily align with our body’s actual physical borders (Ataria, 2014). While the latter are defined by their anatomical structure, the former can fluctuate widely, ranging from feeling strongly separated from the environment to experiencing an almost seamless continuity with it (Ataria, 2014; Ataria et al., 2015; Dambrun, 2016). To date, body boundaries have primarily been investigated in terms of their experienced sharpness, whereas the accuracy with which this boundary can be located in space remains poorly understood. Here, we focus on body boundaries as the perceptual edge of the anatomical body; that is, the precise location of where our physical body ends.

Perceiving the boundaries of our body likely relies on underlying body representations, which broadly refers to perceptual, cognitive, sensorimotor, and affective processes that shape our experience of bodily form, position, movement, and ownership (Gallagher, 2006). While it is widely acknowledged that multiple interacting systems (e.g., body schema, body image) contribute to this representation, there is still active debate regarding the precise number and specific functions of these components (de Vignemont, 2010; Medina and Coslett, 2010; Riva, 2018). At the very least, recent frameworks distinguish between two levels of body representation: somatoperception and somatorepresentation (Longo et al., 2010). Somatoperception refers to real-time, sensory-driven processes, such as tactile localization, proprioceptive tracking, and immediate judgments of body dimensions, whereas somatorep-resentation encompasses more stable, higher-order beliefs about one’s body, such as semantic knowledge of body-part arrangement.

So far, research on somatoperception has primarily focused on coarse, large-scale representations (e.g., entire limbs). These studies have consistently revealed substantial and systematic distortions in body representations (see Longo, 2022, for a comprehensive review). For instance, investigations into perceived body part size have demonstrated significant inaccuracies in judging their relative dimensions (Linkenauger et al., 2015), volume (Sadibolova et al., 2019), or when localizing landmarks (Longo and Haggard, 2010; Mora et al., 2018; Myga et al., 2021). Although these studies have significantly advanced our understanding of body representation, the precision with which fine-grained local body geometry is represented remains largely unexplored. Research on tactile distance perception (Longo and Haggard, 2011) has partially addressed this gap, revealing smaller scale perceptual distortions that reflect biases in the cortical somatosensory homuncular map (Miller et al., 2016). However, these investigations typically consider body parts as mostly flat, two-dimensional surfaces, neglecting its inherently three-dimensional structure. Even seemingly flat regions, such as a hand resting on a table, have pronounced depth and curvature and exhibit intricate three-dimensional geometric structures.

Here, we address this gap by examining whether people can accurately judge their own three-dimensional body boundaries at a fine-grained scale across five individual experiments. First, we compare boundary perception in two body regions with contrasting sensory profiles: the hand—a highly visible, tactilely sensitive, and frequently used body part—versus the ankle, which is typically out of sight and less sensitive. Next, we investigate how changing hand posture might influence boundary judgments, thereby testing whether the underlying representations are sufficiently flexible to adapt to shifts in proprioceptive feedback. We then ask whether a detailed three-dimensional body model is required to explain people’s performance or whether coarser representations suffice. Finally, we investigate whether participants whose body deviates from the average shape make judgements aligned with their own body geometry.

## Results

We developed a novel psychophysical paradigm for assessing the accuracy of the perception of the body boundary. We were specifically interested in quantifying the extent to which the local three-dimensional geometry of different body parts is perceptually accessible in the absence of vision. Participants received simultaneous touches at two predefined points on a given body part and judged whether the midpoint between these points fell within or outside their body. For instance, the midpoint connecting points G and F might objectively fall outside the boundary of the hand (Figure 1A,B, top), while the midpoint connecting points C and E might fall inside (Figure 1A,B, pink). Across experiments, we defined 13 such point pairs spanning inside, outside, and near-boundary locations. To account for individual anatomical differences, we recorded 3D scans of each participant’s body part, then calculated the actual distance of each midpoint from the true boundary. Finally, combining these distances with each participant’s responses (across seven randomized trials per midpoint) enabled us to fit psychophysical curves and estimate both bias and boundary width (Figure 1C).

**Figure 1.**
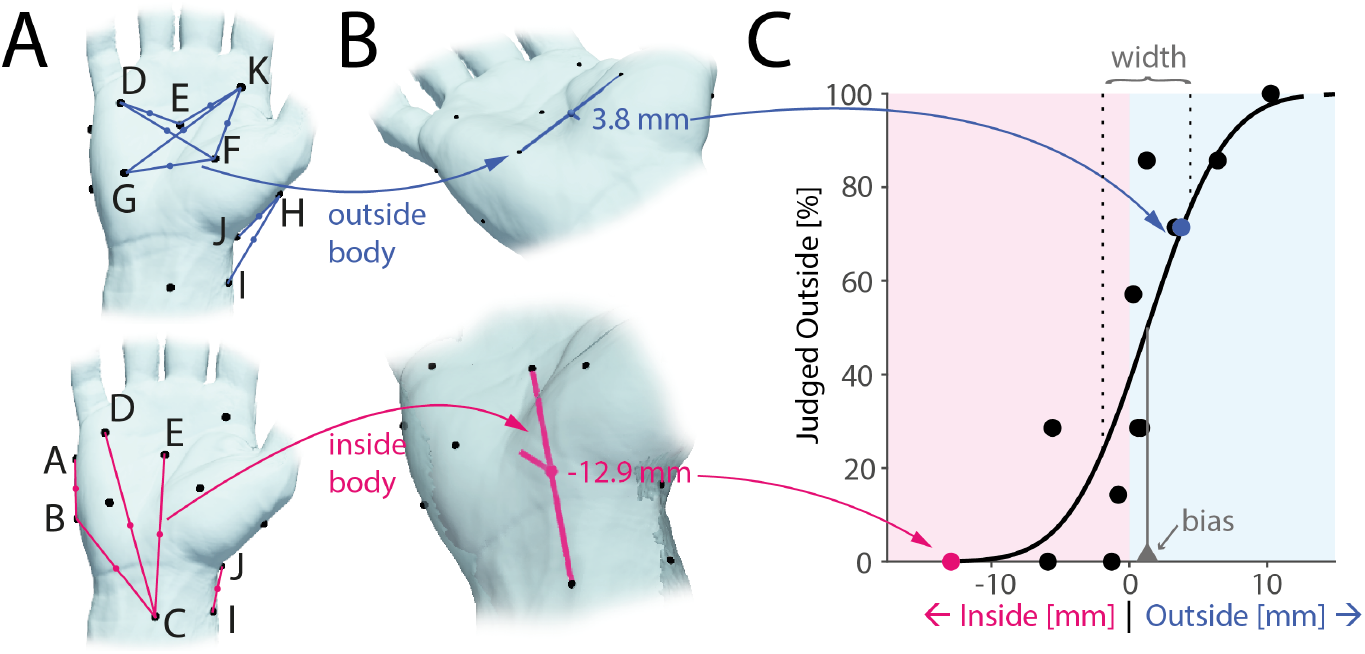
Experimental paradigm. **A**.Participants were touched simultaneously at two points (black dots) and asked to judge whether the midpoint of the imaginary line connecting both touches (colored lines and dots) was perceived to lie inside or outside of their body. Top: Midpoints objectively outside the body for a single participant shown in blue. Bottom: Mid-points inside the body for the same participant shown in pink. **B**. 3D scanning was used to establish the true distance of each midpoint from the body boundary (two examples shown). **C**. A psychophysical curve (black line) was fit to the perceptual judgments (dots) allowing quantification of perceptual bias (vertical line at 50% perceptual threshold) and width of the body boundary (space between vertical dashed lines at 25% and 75%, respectively).

### Millimeter accuracy of body boundary perception on the hand

We first examined participants’ ability to locate their right hand’s boundary while it rested in a relaxed, relatively flat position on a table. Across two separate experiments, each using a different set of midpoints (see Figure 2A) and including 39 participants in total, our results indicate that most participants (29 out of 39, 74%) performed significantly better than chance (as assessed by one-sample chi-squared tests, p < 0.05), many of which were remarkably accurate as demonstrated by steep and well-fit psychophysical curves (see Figure 2B for examples).

**Figure 2.**
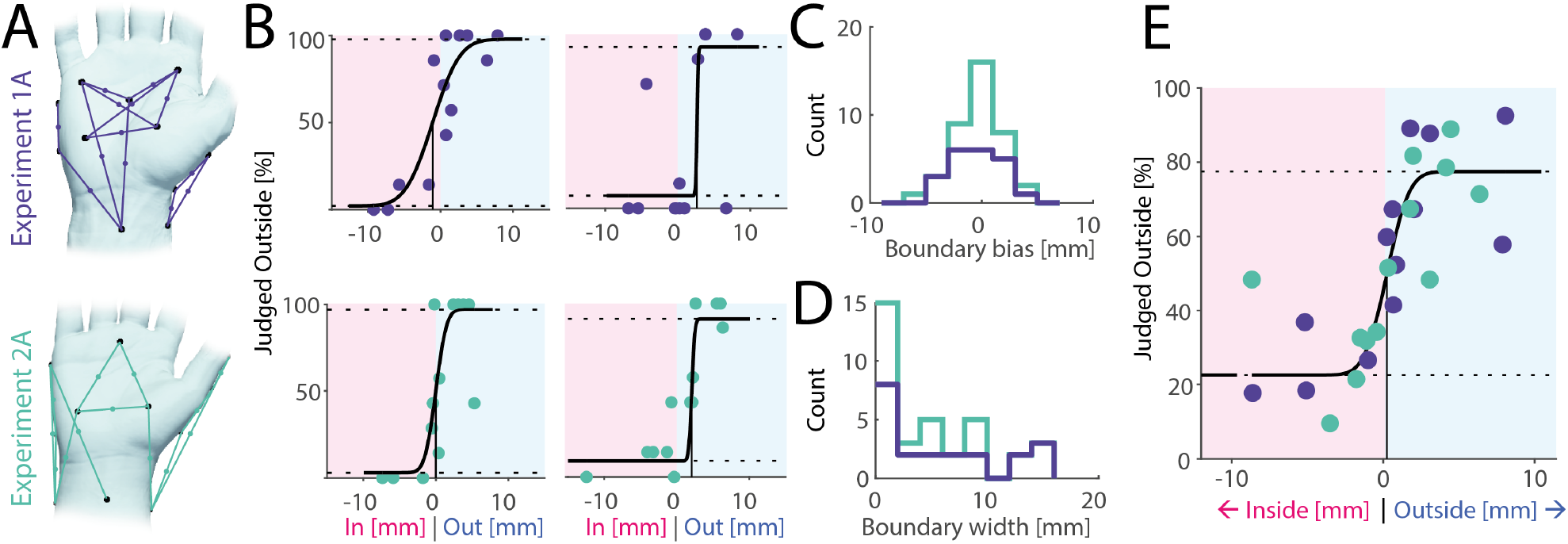
Body boundary perception on the hand. **A**.Touch point pairs and their corresponding midpoints tested in Experiment 1A (n = 21, top) and Experiment 2A (n = 18, bottom) illustrated for a single participant each. In both experiments, the hand was in a relaxed position, resting relatively flat on a table. **B**. Examples of individual fitted psychophysical curves from both experiments. **C**. Perceptual bias (point of subjective equivalence) across all participants. Bias was close to zero and the subjective body boundary was as likely to fall slightly outside the true boundary as it was to fall inside. **D**. Perceptual width of body boundary as assessed by the distance between judging midpoints as mostly inside (25% threshold) and outside (75% threshold). Almost half of the participants showed accuracy within a single millimeter and almost all within one centimeter. **E**. Group-level psychophysical curve across both studies averaging across individual midpoint distances and responses across 26 unique midpoints.

Across all participants, the bias in boundary perception (assessed as the point of subjective equivalence) was on average −0.4 mm (see Figure 2C for a histogram), not significantly different from zero (t(38) = −1.17, p = 0.25, one-sample t-test) and did not differ between the two studies (t(37) = −1.1, p = 0.28, two-sample t-test). Thus, there is no inherent bias to perceive the body boundary as either concave or convex. To assess the width of the body boundary, we calculated the change in distance necessary to jump from a 25% likelihood that a stimulus was judged as outside to a 75% likelihood. On average, the boundary width was 4.9 mm, though for almost half of the participants it was less than 2 mm (Figure 2D). Thus, decisions on whether individual points in space lie inside or outside of the body are made accurately within a few millimeters and often within a single millimeter.

Lastly, by combining data from both experiments and fitting a single psychophysical curve at the group level (Figure 2E), we again confirmed millimeter-level accuracy without systematic bias. Nonetheless, some individual points clearly stood out as consistently misclassified by multiple participants, despite their anatomical position being unambiguously inside or outside the body (see examples in Figure 2B, top right and bottom left). These recurring errors highlight localized perceptual anomalies rather than random judgment mistakes.

### High fidelity of body boundary perception on the ankle

Next, we tested whether body boundary perception would differ on another body part, the ankle, which similar to the hand exhibits intricate local variations in 3D geometry amenable to testing by our paradigm. Across two experiments (2A: n = 21; 2B: n = 7), we collected perceptual judgements across 13 different paired touches, following the same procedure established for the hand (see Figure 3A for touch points and Figure 3B for pairs and respective midpoints). The average bias in boundary perception was 0.6 mm (see Figure 3C for histogram), not significantly different from zero (t(27) = 1.66, p = 0.11, one-sample t-test) or from the bias for the hand (t(65) = 1.95, p = 0.06, two-sample t-test). The boundary width, at an average of 7.3 mm (Figure 3D), was larger than for the hand, though this difference was also not significant (t(61) = 1.68, p = 0.10). These trends were confirmed in the group-level psychophysical curve (Figure 3E), which displayed no bias and high accuracy across 26 midpoints. In contrast to the results from the hand, no clear outliers were present on the ankle. Thus, boundary perception is roughly equally accurate on the ankle compared to the hand.

**Figure 3.**
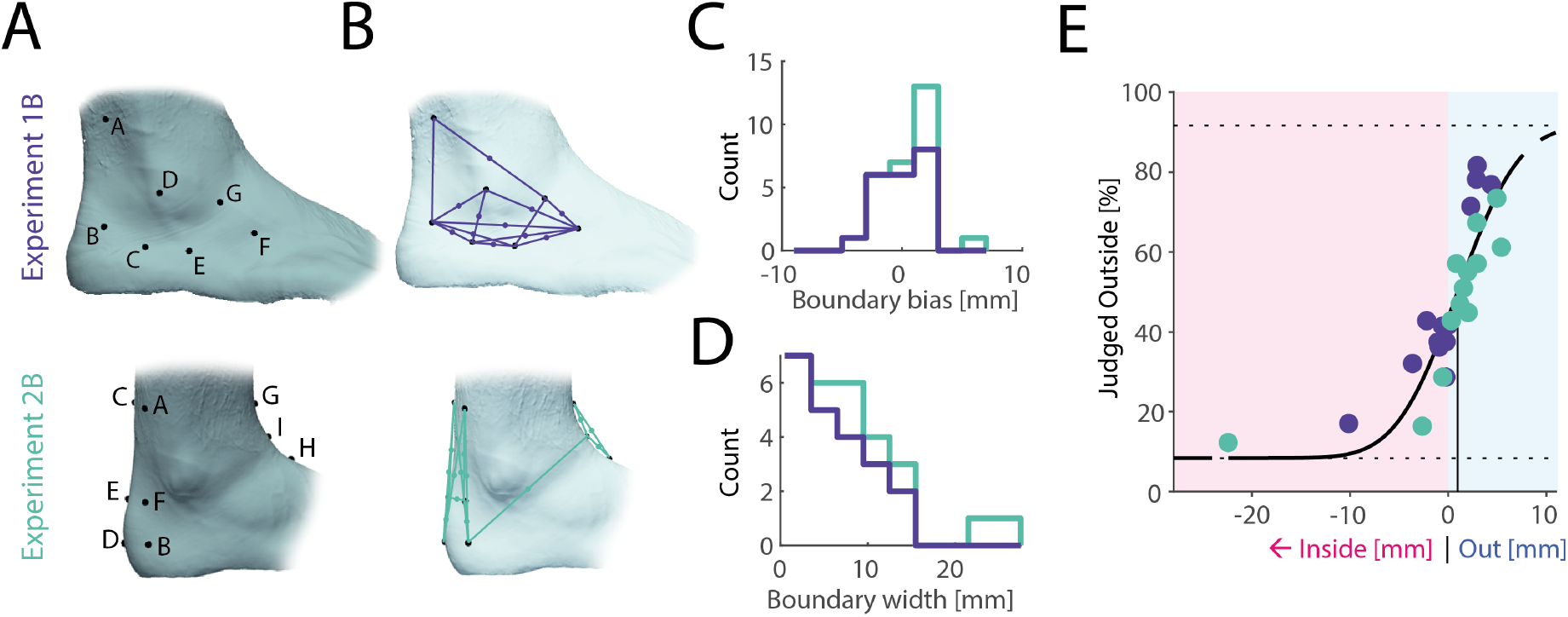
Body boundary perception at the ankle. **A**.Touch points on the ankle for Experiment 1B (n = 21) and Experiment 2B (n = 7). **B**. Corresponding touch point pairs and respective midpoints, focusing on the local geometry around the lateral malleolus for Experiment 1B, and the anterior and posterior foot geometry around the ankle for Experiment 2B. **C**. Bias across all participants. Analogous to the hand, bias was close to zero. **D**. Boundary width on the ankle for all participants (see Figure 2D for comparison with the hand). Most participants showed accuracy within one centimeter, with accuracy slightly lower than on the hand. **E**. Group-level psychophysical curve across both studies averaging across individual midpoint distances and responses across 26 unique midpoints.

### Boundary perception adjusts with changes in posture

A clear difference between the hand and the ankle is the high mobility of the hand with many degrees of freedom. Specifically, different hand postures can lead to drastic differences in whether midpoints fall inside or outside the body. To determine whether hand posture is taken into account when making perceptual body boundary judgments, we tested the same 13 midpoints under two different postures in a subset of participants (n = 11): the baseline relaxed posture and a more “unnatural” posture involving ulnar deviation at the wrist and partial flexion of the thumb (Figure 4A). This second posture induced shifts in the tested midpoints, such that some moved from inside the body to outside and vice versa: midpoints on the ulnar side of the hand moved towards the outside due to ulnar deviation, while midpoints on the hand’s radial side tended to move inside due to flexion of the thumb (Figure 4B). We found that participants’ perceptual judgments shifted in parallel, matching these positional changes in 11 out of 13 mid-points (Figure 4C), and the magnitude of perceptual shift was strongly correlated with the physical shift (r = 0.87, Figure 4D).

**Figure 4.**
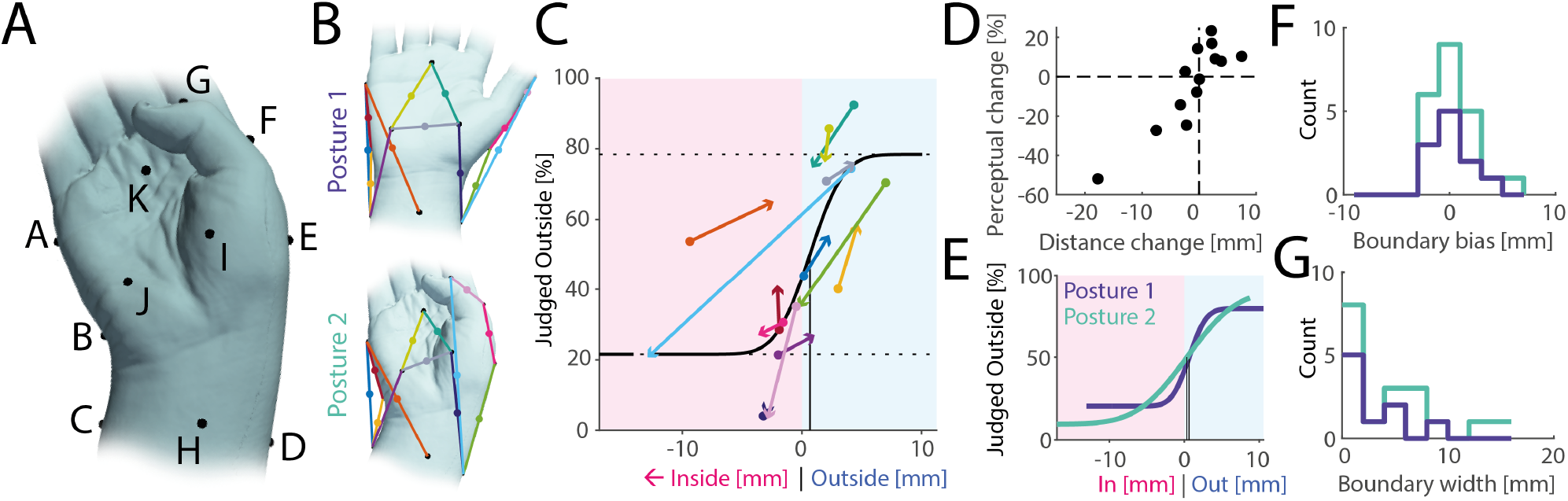
Boundary perception adjusts with changes in posture. **A**.Hand posture adopted in Experiment 2C, characterized by ulnar deviation and partial flexion of the thumb. **B**. Posture 2 was chosen such that, compared to the original baseline posture 1, many midpoints moved with respect to the body boundary, transitioning from inside to outside and vice versa. **C**. Group-level psychophysical curve (black line) along with boundary distances and perceptual judgements averaged over participants: each coloured line denotes a tested midpoint (same colours as in panel B), with round symbols denoting data from posture 1 and arrow heads denoting data from posture 2. As midpoints move from inside the body to outside, perceptual judgements do the same, and vice versa. **D**. Change in distance of midpoints from the body surface between postures 1 and 2 compared to accompanying perceptual change, averaged over all participants. Positive distance change values indicate points moving towards the outside of the body, while negative values denote points moving towards the inside. Positive perceptual change values indicate a higher percentage of stimuli being judged as outside, and vice versa. **E**. Group-level psychophysical curves for posture 1 (dark blue) and posture 2 (green). Their bias is identical and close to zero. The curve is slightly steeper for posture 1. **F**. Individual estimates for bias in boundary perception for both postures. Bias is on average close to zero and does not differ between postures. **G**. Boundary width for all participants across both postures. The width does not differ between postures.

At the group level, the psychophysical curves for both postures were highly similar (Figure 4E). Although the slope was slightly shallower for the second posture, the difference was not statistically significant due to overlapping confidence intervals and no difference in bias. Individual-level analyses also showed no significant differences in bias (t(10) = −0.40, p = 0.70, paired t-test, Figure 4F) or boundary width (t(7) = −1.49, p = 0.18, paired t-test, Figure 4G) between postures. We conclude that body boundary perception reflects changes to posture, maintaining high accuracy.

### Boundary perception reflects local three-dimensional body geometry

We next investigated whether participants’ judgments depended on detailed three-dimensional body representations or on simpler, two-dimensional outlines influenced by posture alone. To do this, we progressively smoothed each 3D hand model (Figure 5A, bottom row) and then compared participants’ judgments with the new, smoothed boundaries (see Methods). If people rely on fine-grained local geometry, removing these details should reduce accuracy. Conversely, if the mental representation is relatively coarse, smoothing might not affect performance; or could even improve it. Since smoothing does not change the broad outline of the hand, but instead affects local geometry only, we divided all mid-points tested in the hand baseline condition across both exper-iments into two categories: those reflecting predominantly local 3D geometry (see coloured markers in Figure 5B, located on the palm) and those reflecting simple 2D body outlines or the coarse postural configuration of the hand (grey markers in Figure 5B). For example, the palm area exhibits local to-pography even when the hand is resting flat on the table and correct judgements would therefore require a body representation incorporating the shape and curvature of the thenar and hypothenar eminences. In contrast, other judgements, such as ones about the line connecting the metacarpophalangeal joint of the thumb and the wrist, might be resolved using a simpler 2D representation of the hand combined with postural feedback.

**Figure 5.**
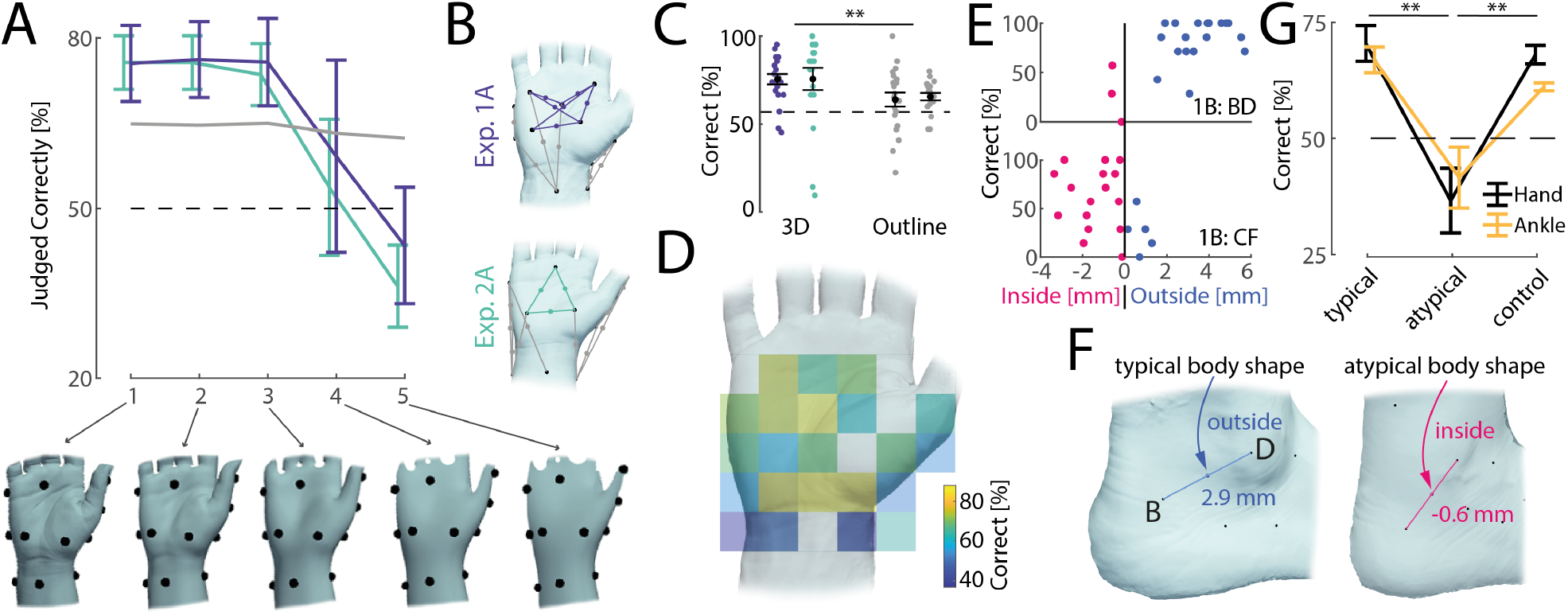
Boundary perception reflects local, typical geometry of the body. **A**.Accuracy of perceptual judgements when recalculated with respect to progressively smoothed models of the hand (1: measured high-resolution model, 5: heavily smoothed), shown below. Coloured traces refer to perceptual judgements related to local 3D structure of the hand (as shown in panel B), with lines reflecting averages and error bars denoting standard error of the mean across different midpoints. The grey line is the average accuracy for all other perceptual judgments, mostly reflecting the 2D outline of the hand (see panel B). The dashed line indicates chance level. Accuracy drops with any amount of smoothing, but worsens considerably for models 4 and 5. **B**. Midpoints included in smoothing analysis shown in panel A using the same colour scheme. **C**. Accuracy of perceptual judgements reflecting local 3D structure (left) versus 2D hand outlines (right). Black markers denote the average and standard error of the mean across participants, with individual data points shown as coloured dots (same colour scheme as in panel B). **D**. Accuracy of perceptual judgements when plotted spatially according to the location of the judged midpoint. Each judgment from every participant (n = 39) in the hand baseline posture (Experiments 1A and 2A) was first mapped onto the reference hand shown using procrustes analysis, after which averages were calculated for each spatial bin. Accuracy is markedly lower just below the wrist and on both sides of the hand, but high on the palm. **E**. Examples of midpoints on the ankle (Experiment 1B) falling on different sides of the body boundary for different participants and their associated perceptual judgments. Shown is distance to the actual body boundary on the horizontal axis against the percentage of correct answers. Each point shown is a different participant. Participants whose body boundary does not agree with the typical body shape display low accuracy. **F**. Examples of a typical (left) and atypical (right) body shape on the ankle (taken from Experiment 1B). Typically the lateral malleolus is pronounced enough to cause the midpoint between B and D to fall outside the body boundary. However, for a minority of participants, this body region is flattened instead, leading for the midpoint to fall inside the body. **G**. Average accuracy for hand (black lines, experiments 1A and 2A) and ankle (yellow, experiments 1B and 2B) judgments where the participant’s body shape agreed with the average body shape (“typical”, left), where the participant’s body outline disagreed with the average body shape (“atypical”, middle), and all other judgments made by the same participants included in the atypical dataset (“control”, right). Participants with atypical body shape show much lower accuracy on these judgements than participants with a typical body shape, even though other judgements by these participants do not display lower accuracy.

We found that low amounts of smoothing did not impair participants’ accuracy, but there was a steep decline for more extensive smoothing levels (see coloured lines in Figure 5A), suggesting that local geometry at a high level of detail is both perceptually accessible and important for solving the task (see model 3 in Figure 5A, bottom row, for the smoothest model where accuracy was above chance level). In contrast, since smoothing did not affect the general outline of the hand model or its overall posture, the accuracy of these judgments did not change (see grey line in Figure 5A). Moreover, judgements associated with local 3D geometry were on average more accurate than those merely requiring a 2D outline model of the hand (t(38) = 2.86, p < 0.01, paired t-test, Figure 5C), despite the latter both intuitively appearing easier and objectively being located further from the body surface (mean absolute distance 3.8 mm vs 2.8 mm for 3D midpoints, t(38) = −7.39, p < 0.01, paired t-test). Additionally, a lateralized effect emerged: in Experiment 2A, touch points were balanced such that five judgments included the left side of the hand and five the right side (with three focusing on the palm), and comparing their respective accuracy showed significantly higher performance on the right (73%) compared to the left side (59%, t(178) = −3.25, p < 0.01, two-sample t-test). Indeed, mapping all judgments across both experiments onto a standardized reference hand revealed clear spatial clustering: high accuracy on the palm and consistent errors on the wrist, and the sides of the hand (Figure 5D). That errors are spatially clustered suggest that they arise from underlying distortions in local body representation rather than random lapses in judgment.

### Boundary judgements are aligned with a typical body model

We selected touch points such that their midpoints were predictably inside or outside the body boundary for most participants. However, as this was not always the case, we 3D-scanned the tested body parts for each participant to measure their individual body shape. We found that for 13 out of 26 total midpoints tested on the hand (across Experiments 1A and 2A), some participants’ body shape deviated enough for these points to cross the body boundary from inside to outside or vice versa. Similarly, 11 out of 26 midpoints tested on the ankle (across experiments 1B and 2B) spanned both sides of the body boundary across all participants tested. For example, the lateral malleolus typically presents as a pronounced bump on the ankle, causing the line connecting it to the back of the foot to lie outside the body for most participants (see Figure 5F, left). However, for a small subset of participants this bump was much less pronounced and the surrounding region correspondingly almost flat, such that the line fell inside the body instead (see Figure 5F, right). Are these individual differences in body geometry reflected in participants’ judgments? We noticed that participants’ accuracy was often low for judgements in which their individual body shape differed from the typical one (see examples in Figure 5E). Indeed, when probing this effect across all data points in the hand baseline (Experiments 1A and 2A) and the ankle (Experiments 1B and 2B) conditions, we found that perceptual judgments were on average much more accurate in cases when the participant’s body shape agreed with the average shape compared to when the participant’s body differed from the average (Figure 5G, typical vs atypical columns). This effect could not be explained by participants with an atypical body exhibiting lower accuracy generally, as their other judgments for different midpoints were much more accurate (Figure 5G, control column). In line with these observations, an ANOVA found significant differences between typical, atypical and control judgments (F(2,52) = 11.16, p < 0.01) and post-hoc Tukey tests confirmed that atypical judgements differed from the two other classes (p < 0.01 for both), while the accuracy of typical and control judgments did not differ (p = 0.88). We note that the difference between typical and atypical body shapes was small, with the body boundary shifting by only 2.1 mm on average between typical and atypical body shapes. Larger discrepancies might well be integrated into people’s body representations. However, on a small scale, when there are individual differences in body shape, participants appear to align their judgments with a more ‘typical’ body rather than their own.

## Discussion

We developed a novel psychophysical paradigm to examine how precisely people judge the boundaries of their own body, specifically whether the midpoint between two brief touches on the skin fell inside or outside the body. Across five experiments, we systematically tested boundary judgments in three conditions: the hand in a relaxed posture (Experiments 1A and 2A), the hand in an “unnatural” posture (Experiment 2C), and the ankle (Experiments 1B and 2B). Despite differing sensory profiles and visual familiarity across these body parts, our findings consistently showed that participants could localize their body boundaries with often striking accuracy—within just a few millimeters of the actual anatomical border—even without visual feedback and when assuming different postures.

### Comparison with previous findings

Previous work on body representations has emphasized large perceptual distortions, which replicate robustly and transfer across different experimental protocols (Longo, 2022). How can our finding of high perceptual accuracy be reconciled with these prior results? First, the apparent discrepancy might be a consequence of our experimental paradigm. Previous studies, whether on body part size estimation or judgements about tactile distance, specifically asked participants to make size or length estimates. A distorted representation of the hand, for example stretched or compressed along a certain axis, is easily measurable using these paradigms. In contrast, our paradigm assessed whether people are aware of the shape of their body between two touched points. Any systematic distortion (stretch or compression) of these body parts would not affect their overall shape, and perceptual judgments should therefore be unaffected. Thus, if body representations are fine-grained and accurate in their three-dimensional representation, but otherwise distorted by appearing elongated or truncated along a certain axis, then our findings are entirely compatible with previous ones. Second, it is possible that we tested body parts where representations are relatively undistorted. The large distortion of the hand representation is mainly evident on the hand dorsum and considerably smaller on the palmar side (Longo and Haggard, 2012). While perceptual distortions have been measured on the foot dorsum (Manser-Smith et al., 2021), to our knowledge the ankle has not yet been tested specifically. Additionally, recent findings suggest that observed distortions might be a consequence of cognitive processes rather than the underlying body representation (Peviani et al., 2024). Our findings demonstrate that local 3D body geometry is accurately represented. Finally, it is possible that our task, specifically asking participants to make judgements about their body boundaries, engaged different representations than those in previous tasks. Given the robustness of previous findings with respect to the specific paradigm used, we believe this to be unlikely.

### Regional differences in body boundary perception

Our results indicated that while the width of the perceived body boundary was larger on the ankle compared to the hand, this difference was not significant and, in any case, smaller than we had anticipated. We predicted a larger discrepancy because the hand possesses denser mechanoreceptor innervation (Corniani and Saal, 2020), better tactile spatial discrimination and localization (Weinstein, 1968; Mancini et al., 2014), lower tactile thresholds (Hennig and Sterzing, 2009), a disproportionately large cortical representation (Hennig and Sterzing, 2009), and higher proprioceptive accuracy (Han et al., 2013) compared to the ankle. Moreover, our hands frequently occur within our field of view during manual tasks (Mineiro and Buckingham, 2023), and the space close to our hands draws attention towards it (Reed et al., 2006), which is not the case for the foot. Previous studies have noted the importance of vision in shaping body representations (Rakesh Kottu and Lazar, 2024; Shahzad et al., 2025). Nevertheless, body boundary perception was similar on both body parts, which appear to be represented with high fidelity.

While there was no significant difference between body boundary perception on the hand and the ankle, perceptual accuracy differed spatially across the hand. Specifically, error rates were high on the wrist, even when judging locations that were clearly inside or outside the body. In fact, there appeared to be a sharp edge in accuracy between the hand and the wrist, with the lower palm area judged highly accurately, while the area just proximal to the wrist border was much less so. Similar localized errors have been reported in prior studies of tactile distance perception, where distortions can cluster around joint boundaries or transitional skin areas, possibly because of segmented body representations (de Vignemont et al., 2009; Knight et al., 2014; Tamè and Longo, 2023). A similar effect might be responsible for our findings.

### Fine-grained representation of local body geometry

We found that body boundary perception adapted to changes in posture, meaning that ongoing feedback about the body’s configuration is therefore taken into account when making such judgements. We did not find significant differences in boundary perception between the two postures tested, but previous literature has described postural effects on other measures of body representation (Longo, 2015). Importantly, postural information alone could not explain the observed results, as coarsening the measured 3D body part models led to considerable decreases in accuracy. In fact, somewhat counterintuitively, we found that participants were significantly more accurate when judgements involved detailed 3D body shape, rather than two-dimensional outlines. These results suggest that participants’ body models incorporate detailed three-dimensional shape information. In the absence of vision, somatosensory feedback might supply some relevant information: skin stretch patterns contain information about three-dimensional body conformation beyond that supplied by proprioceptive feedback (Rupani et al., 2025), and some tactile neurons in the hand have been found to provide ongoing information about the local state of the skin even in the absence of an external force (Saal et al., 2023). In line with these findings, anesthetizing body parts has been found to change perceived body geometry (Melzack and Bromage, 1973; Inui et al., 2011; Walsh et al., 2015), suggesting that ongoing feedback is required to maintain these representations and perhaps contributes to them (see also Crucianelli and Ehrsson, 2023, for somatosensory contributions to interoception more broadly). Nevertheless, body models are likely to be constructed and stored, as it is unlikely that all relevant information would be available from sensory feedback (Longo and Haggard, 2010). Our finding that in some cases participants’ judgements aligned with a typical rather than their own body would appear to support this statement.

### Boundary perception might reflect typical rather than individual geometry

We found that when a midpoint fell inside or outside the body boundary for most participants, even those whose anatomy diverged from this norm often judged it in line with the majority’s boundary. This suggests that body boundary perception is not determined solely by real-time sensory input, but also by a generalized, internalized model of the body (Haggard et al., 2003; Longo and Haggard, 2010; Tsakiris, 2010). One possible explanation for this effect involves statistical learning: over time, individuals observe countless bodies (both their own and others’), gradually internalizing a generalized prototype of where their boundaries should lie. Consequently, when deprived of visual feedback, participants may default to this prototype rather than rely on their unique anatomy. Supporting this idea, previous research has found that people often fail to recognize photographs of their own hands (Wuillemin and Richardson, 1982; Holmes et al., 2022) and that weaker visual memories of one’s body correlates with heightened plasticity in its representation (O’Dowd and Newell, 2020), highlighting how higher-order body models are not fully individualized and can sometimes override immediate sensory cues.

Notably, differences between typical and atypical body shapes were relatively subtle in our study. An interesting question for future research is to what extent larger deviations in body shape are integrated into individual body models, which could be tested by combining our paradigm with the rubber hand illusion or manipulation of visual feedback in virtual reality. Testing other body regions such as the neck, elbow, or other joints could also help determine whether observed perceptual patterns hold true throughout the body or vary systematically according to anatomical or functional characteristics. The face in particular represents an intriguing target due to its unique and individualized anatomical features, potentially challenging participants’ reliance on generalized, socially-derived body representations. Finally, future work should also test representations across body segments to examine the origin of systematic and anatomically incorrect judgments. For example, many participants judged midpoints near the wrist inaccurately, even if these judgments clearly defied plausible anatomical limits. An “average” body model would not reliably mislabel these stimuli. Instead, these rare but systematic inaccuracies indicate the presence of localized distortions or other processing issues within the underlying body representation that warrants further investigation.

In summary, we found that detailed three-dimensional body geometry is perceptually accessible, with high accuracy both on the palmar surface of the hand and on the ankle. Perceptual judgments of the body boundary adapted to changes in posture, maintaining accuracy and demonstrating the integration of ongoing sensory feedback. Judgments requiring knowledge of detailed, local geometry displayed the highest accuracy, and detailed local geometry is therefore represented, though this internal model might partially rely on general knowledge of the shape of the typical body, rather than participants’ own.

## Methods

### Participants

Twenty-one individuals (twelve female, nine male) between the ages of 18 and 27 participated in Study 1. All participants were right-handed, except for one who was ambidextrous. Study 1 was composed of two experiments: psychophysical testing on the hand (Experiment 1A), followed by the ankle (Experiment 1B). All twenty-one participants completed both experiments. Eighteen participants (thirteen female, five male) aged 18 to 28 participated in study 2. All but one participant was right-handed. Study 2 consisted of three experiments. The first experiment (Experiment 2A), testing the hand using new touch points, was completed by all participants. Seven of them subsequently completed the second experiment (Experiment 2B), which included points on the ankle, while the remaining eleven participants completed the third experiment, which tested the hand again, but in a different posture (Experiment 2C). All participants provided written informed consent prior to the start of data collection. The study protocol was approved by the ethical review board of the School of Psychology at the University of Sheffield (protocol number 060858).

### Selection of touch points, midpoints, and postures

#### Experiments 1A and 1B

To ensure uniform coverage of perceptual midpoints across participants, predefined locations were selected on the hand and ankle. The chosen points were selected to maximize variation in midpoint locations, ensuring that midpoints were either robustly and clearly falling inside the body, outside of it, or very close to the surface. For the hand (Experiment 1A), participants were instructed to rest their right hand in a relaxed posture on the table. Eleven touch points (labeled A–K) were identified on the palmar surface of the hand and the wrist area. Thirteen unique pairs of points were selected to generate midpoints spanning the inside, outside, and boundary of the hand. For the ankle (Experiment 1B), participants were seated in a chair, resting their right foot on the ground at a roughly right angle. Seven touch points (labeled A through G) around the lateral malleolus were identified and then 13 pairs were chosen such that their midpoints again spanned the inside and outside of the body in a range similar to that of the hand (Figure 3A,B, top row).

#### Experiments 2A, 2B, and 2C

In Experiment 2A, the hand was in the same posture as in Experiment 1A. For Experiment 2C, participants adopted a new posture, characterized by ulnar deviation at the wrist and partial flexion of the thumb (see Figure 4A). This posture was chosen to maximize changes in the distance of the midpoints to the body surface between the two postures. The same set of touch points and midpoints were tested in both experiments. A new set of 11 points (labeled A through K) was chosen, with some points identical to those in 1A and others newly chosen. In addition to replicating the results of Experiment 1A with new midpoints, we also wanted to specifically force perceptual judgments to be either about boundary depth (i.e. local 3D geometry) or two-dimensional outlines (e.g. edge judgments). The hand was chosen for the posture experiment, because of its high mobility with many degrees of freedom. For Experiment 2B, we focused on the ankle again, but chose touch points further away from the lateral malleolus, testing the foot outline around the ankle more broadly, rather than detailed local geometry. A set of 9 touch points (labeled A-I) was chosen, from which again 13 pairs were selected (Figure 3A,B, bottom row).

### Experimental Procedure

Participants were blindfolded throughout the experiment to eliminate visual feedback. A skin-safe marker pen was used to label the predefined touch points on the skin. After labeling, the participant assumed the tested posture, for which they received gentle guidance by the experimenter, if needed. The relevant body part (hand or ankle) was then 3D scanned. Scanning was performed using a POP 3 Plus (Revopoint 3D, Shenzhen, China) handheld 3D scanner and the accompanying RevoScan software, which generated 3D models. Scanning took a couple of minutes, after which the psychophysical experiment started.

For each individual trial, participants received brief simultaneous touches at two predefined touch points using metal probes with rounded tips roughly 2 mm in diameter. Touches were light and consisted of brief taps on the skin. These were performed by the same experimenter for all participants. Participants were required to make a two-alternative forced choice judgment on whether the perceived midpoint between the two touches fell inside or outside of their body boundary and noted by an experimenter. Each experiment included 13 pairs of touch points, which were tested 7 times each, for a total of 91 trials per experiment. The order of stimuli was randomized to prevent response bias and order effects. In the first trial, participants were instructed to take as much time as needed to respond. For subsequent trials, participants had up to 3 seconds to provide a response. Participants were allowed to take breaks or to relax their hands for a few seconds in between trials, but the majority completed the full protocol in one go. If a pause was taken, participants were carefully guided by the experimenter to assume the original posture. In study 1, participants first completed trials on the hand before moving on to the ankle. In study 2, participants first completed trials in the relaxed hand posture, before moving on to either the ankle (first 7 participants) or the second hand posture (remaining 11 participants).

### Analysis

#### 3D model processing

Using custom code based on the *pyvista* Python module, touch points were manually identified from the pen markings on the obtained 3D models. Midpoints and their distances from the skin surface were then calculated and exported as csv files.

#### Psychophysical Curve fitting

Psychophysical curves were fitted to the data using the *psignifit* Matlab package (Schütt et al., 2016). Cumulative Gaussian sigmoids were chosen as psychophysical functions together with a single guess rate, as there was no reason to assume any difference in error rates between inside and outside choices. For group-level psychophysical curves, we first averaged distances and perceptual judgments across participants for each midpoint, before fitting a single psychophysical curve. Thresholds at 25%, 50%, and 75% were calculated from the psychophysical curves. The 50% threshold was taken as the bias. Width of the body boundary was defined as the physical distance between the 25% and 75% thresholds. Measures of fit differed across participants, but all obtained data was included in all further analyses.

#### Smoothing analysis

To test whether detailed three-dimensional body shape was taken into account by participants, we calculated whether their judgements were accurate with respect to a series of progressively smoothed body part models. For smoothing, we used the *smooth-SurfaceMesh* function, part of the Lidar Toolbox for Matlab 2024a (Mathworks, Natick, MA), which implements a simple averaging filter that iteratively computes a local weighted mean average on a given mesh. This function was run for 50, 500, 2500, and 5000 steps to yield smoothed models 2-5, respectively, with model 1 being the original unsmoothed model. This analysis was run for all hand models in the relaxed posture (Experiments 1A and 2A, n = 39). We then recalculated the location of each midpoint and its distance to the surface for each of the smoothed models. Finally, we checked whether participants’ responses were correct with respect to these new distances. This analysis was run for each individual participant separately, before averaging.

#### Comparing accuracy for typical and atypical body shapes

To test whether participants whose body shape differed from the typical body aligned their judgments with their own body or the average, we first determined all midpoints which fell on different sides of the body boundary for at least one participant across all data from the hand in the relaxed posture (Experiments 1A and 2A, n = 39) and the ankle (Experiments 1B and 2B, n = 28). For each of these we then calculated the average distance and took the side of the body boundary on which it fell (inside or outside) as the ‘typical’ body shape. We then calculated the average accuracy of all participants who agreed with this average (typical) and the average accuracy of all who disagreed (atypical). If participants with an atypical body exhibit high accuracy, this indicates that they are judging their own body; conversely, if they display low accuracy, their internal body representation might agree more with the typical shape. For each midpoint and atypical participant, we also calculated their accuracy on all other midpoints (control) to test whether these participants generally show worse performance, independent of any anatomical differences in body shape.

## ACKNOWLEDGEMENTS

This work was funded by the Leverhulme Trust under Research Project Grant RPG-2022-031.

## Bibliography

Ataria, Y., Dor-Ziderman, Y., and Berkovich-Ohana, A. How does it feel to lack a sense of boundaries? a case study of a long-term mindfulness meditator. Conscious. Cogn., 37: 133–147, Dec. 2015.

Ataria, Y. Where do we end and where does the world begin? the case of insight meditation. Philos. Psychol., 28(8):1128–1146, Nov. 2014.

Corniani, G. and Saal, H. P. Tactile innervation densities across the whole body. J. Neurophysiol., 124(4):1229–1240, Oct. 2020.

Crucianelli, L. and Ehrsson, H. H. The role of the skin in interoception: A neglected organ? Perspect. Psychol. Sci., 18(1):224–238, Jan. 2023.

Dambrun, M. When the dissolution of perceived body boundaries elicits happiness: The effect of selflessness induced by a body scan meditation. Conscious. Cogn., 46:89–98, Nov. 2016.

de Vignemont, F., Majid, A., Jola, C., and Haggard, P. Segmenting the body into parts: evidence from biases in tactile perception. Q. J. Exp. Psychol., 62(3):500–512, Mar. 2009.

de Vignemont, F. Body schema and body image–pros and cons. Neuropsychologia, 48(3): 669–680, Feb. 2010.

Gallagher, S. How the body shapes the mind. Clarendon Press, Oxford, England, Oct. 2006.

Haggard, P., Taylor-Clarke, M., and Kennett, S. Tactile perception, cortical representation and the bodily self. Curr. Biol., 13(5):R170–3, Mar. 2003.

Han, J., Anson, J., Waddington, G., and Adams, R. Proprioceptive performance of bilateral upper and lower limb joints: side-general and site-specific effects. Exp. Brain Res., 226 (3):313–323, May 2013.

Hennig, E. M. and Sterzing, T. Sensitivity mapping of the human foot: thresholds at 30 skin locations. Foot Ankle Int., 30(10):986–991, Oct. 2009.

Holmes, N. P., Spence, C., and Rossetti, Y. No self-advantage in recognizing photographs of one’s own hand: experimental and meta-analytic evidence. Exp. Brain Res., 240(9): 2221–2233, Sept. 2022.

Ihde, D. Sense and significance. Duquesne University Press, 1973.

Inui, N., Walsh, L. D., Taylor, J. L., and Gandevia, S. C. Dynamic changes in the perceived posture of the hand during ischaemic anaesthesia of the arm. J. Physiol., 589(Pt 23): 5775–5784, Dec. 2011.

Knight, F. L. C., Longo, M. R., and Bremner, A. J. Categorical perception of tactile distance. Cognition, 131(2):254–262, May 2014.

Linkenauger, S. A., Wong, H. Y., Geuss, M., Stefanucci, J. K., McCulloch, K. C., Bülthoff, H. H., Mohler, B. J., and Proffitt, D. R. The perceptual homunculus: the perception of the relative proportions of the human body. J. Exp. Psychol. Gen., 144(1):103–113, Feb. 2015.

Longo, M. R. and Haggard, P. An implicit body representation underlying human position sense. Proc. Natl. Acad. Sci. U. S. A., 107(26):11727–11732, 2010.

Longo, M. R. and Haggard, P. Weber’s illusion and body shape: Anisotropy of tactile size perception on the hand. J. Exp. Psychol. Hum. Percept. Perform., 37(3):720–726, 2011.

Longo, M. R. and Haggard, P. A 2.5-D representation of the human hand. J. Exp. Psychol. Hum. Percept. Perform., 38(1):9–13, 2012.

Longo, M. R., Azañón, E., and Haggard, P. More than skin deep: body representation beyond primary somatosensory cortex. Neuropsychologia, 48(3):655–668, Feb. 2010.

Longo, M. R. Posture modulates implicit hand maps. Conscious. Cogn., 36:96–102, 2015.

Longo, M. R. Distortion of mental body representations. Trends Cogn. Sci., 26(3):241–254, Mar. 2022.

Mancini, F., Bauleo, A., Cole, J., Lui, F., Porro, C. A., Haggard, P., and Iannetti, G. D. Whole-body mapping of spatial acuity for pain and touch. Ann. Neurol., 75(6):917–924, 2014.

Manser-Smith, K., Tamè, L., and Longo, M. R. Tactile distance anisotropy on the feet. Atten. Percept. Psychophys., 83(8):3227–3239, Nov. 2021.

Medina, J. and Coslett, H. B. From maps to form to space: Touch and the body schema. Neuropsychologia, 48(3):645–654, 2010.

Melzack, R. and Bromage, P. R. Experimental phantom limbs. Exp. Neurol., 39(2):261–269, 1973.

Miller, L. E., Longo, M. R., and Saygin, A. P. Mental body representations retain homuncular shape distortions: Evidence from weber’s illusion. Conscious. Cogn., 40:17–25, 2016.

Mineiro, J. and Buckingham, G. O hand, where art thou? mapping hand location across the visual field during common activities. Exp. Brain Res., 241(5):1227–1239, May 2023.

Mora, L., Cowie, D., Banissy, M. J., and Cocchini, G. My true face: Unmasking one’s own face representation. Acta Psychol., 191:63–68, Nov. 2018.

Myga, K. A., Ambroziak, K. B., Tamè, L., Farnè, A., and Longo, M. R. Whole-hand perceptual maps of joint location. Exp. Brain Res., Feb. 2021.

O’Dowd, A. and Newell, F. N. The rubber hand illusion is influenced by self-recognition. Neurosci. Lett., 720(134756):134756, Feb. 2020.

Peviani, V. C., Miller, L. E., and Medendorp, W. P. Biases in hand perception are driven by somatosensory computations, not a distorted hand model. Curr. Biol., May 2024.

Rakesh Kottu, S. and Lazar, L. Lack of visual experience leads to severe distortions in the hand representation of the body model. Cortex, 183:38–52, Oct. 2024.

Ratcliffe, M. Touch and the sense of reality. In Radman, Z., editor, The Hand, an Organ of the Mind: What the Manual Tells the Mental. MIT Press, 2013.

Reed, C. L., Grubb, J. D., and Steele, C. Hands up: attentional prioritization of space near the hand. J. Exp. Psychol. Hum. Percept. Perform., 32(1):166–177, Feb. 2006.

Riva, G. The neuroscience of body memory: From the self through the space to the others. Cortex, 104:241–260, July 2018.

Rupani, M., Cleland, L. D., and Saal, H. P. Local postural changes elicit extensive and diverse skin stretch around joints, on the trunk and the face. J. R. Soc. Interface, 22(223), Feb. 2025.

Saal, H. P., Birznieks, I., and Johansson, R. S. Memory at your fingertips: how viscoelasticity affects tactile neuron signaling. bioRxiv, page 2023.05.15.540820, May 2023.

Sadibolova, R., Ferrè, E. R., Linkenauger, S. A., and Longo, M. R. Distortions of perceived volume and length of body parts. Cortex, 111:74–86, Feb. 2019.

Schütt, H. H., Harmeling, S., Macke, J. H., and Wichmann, F. A. Painfree and accurate bayesian estimation of psychometric functions for (potentially) overdispersed data. Vision Res., 122:105–123, May 2016.

Shahzad, I., Occelli, V., Giraudet, E., Azañón, E., Longo, M. R., Mouraux, A., and Collignon, O. How visual experience shapes body representation. Cognition, 254(105980):105980, Jan. 2025.

Tamè, L. and Longo, M. R. Emerging principles in functional representations of touch. Nature Reviews Psychology, pages 1–13, June 2023.

Tsakiris, M. My body in the brain: a neurocognitive model of body-ownership. Neuropsychologia, 48(3):703–712, Feb. 2010.

Walsh, L. D., Hoad, D., Rothwell, J. C., Gandevia, S. C., and Haggard, P. Anaesthesia changes perceived finger width but not finger length. Exp. Brain Res., 233(6):1761–1771, 2015.

Weinstein, S. Intensive and extensive aspects of tactile sensitivity as a function of body part, sex, and laterality. In Kenshalo, D. R., editor, The skin senses, pages 195–222. Thomas, Springfield, 1968.

Wuillemin, D. and Richardson, B. On the failure to recognize the back of one’s own hand. Perception, 11(1):53–55, 1982.

